# “Double-Effect” strategy against Triple-Negative Breast Cancer *via* a synergistic therapy of Magneto-mechanical Force enhancing NIR-II Hypothermal Ablation

**DOI:** 10.1101/2022.10.03.510118

**Authors:** Hui Du, Fang Yang, Chenyang Yao, Wenhao Lv, Hao Peng, Stefan G. Stanciu, Harald A. Stenmark, Young Min Song, Bo Jiang, Aiguo Wu

## Abstract

Triple-negative breast cancer (TNBC) is a form of breast cancer that is more aggressive and harder to treat than others, with a higher probability of relapse. Its highly efficient capabilities for migrating and invading other parts of the body together with the current lack of clinically established effective therapies account for a low survival rate. Thus, we propose the in-tandem use of two complementary therapeutic routes to effectively combat TNBC. Herein, a versatile magnetic-photothermal converter (MPC) is elaborately designed *via* integrating zinc-doped ferrite nanoparticles and polyethene glycol, synergistically enhanced through magneto-mechanical force (MMF) and near-infrared-II (NIR-II) hypothermal ablation, thereby displaying excellent therapeutic efficiency. Their combined use, which is less aggressive to the human body compared to conventional chemotherapeutic approaches, results in the splendid suppression of TNBC migration and invasion. Remotely controlling the MPCs by an external magnetic stimulus, results in cellular MMF effects that cause direct mechanical destruction to the cancer cell membrane, leading to its necrosis. Furthermore, the MMF disrupts intracellular lysosomes, thereby triggering the release of large amounts of protein hydrolases, which induce intracellular oxidative stress, and accelerate the induction of apoptosis. Complementing the therapeutic approach based on MMF, the excellent photothermal performance of the MPC in the NIR-II region (1064nm) is exploited to enable effective hypothermal ablation of the tumours, which can be achieved in deep tissue layers. The proposed multifunctional nanocomposites, together with the demonstrated “double effect” therapeutic approach, hold significant potential to pave the way for future cutting-edge weapons against the dreadful TNBC.

## 1. Introduction

Triple-negative breast cancer (TNBC) represents a huge threat to human health, and a heavy burden to healthcare systems worldwide, owing to its highly efficient capabilities for migration, invasion and metastasis. The five-year survival rate for patients diagnosed with localized TNBC, not spread beyond the breast, is 91%, but this percentage severely drops to 65% for patients diagnosed at a stage when the TNBC has spread into nearby lymph nodes, and down to 11% for patients in which the TNBC has spread to critical parts of the body, such as bones, lungs or liver [1,2]. TNBC cells are negative for the estrogen receptor (ER), progesterone receptor (PR) and human epidermal growth factor receptor 2 (HER2), thus aren’t fueled by hormones that can be specifically targeted, as in other types of breast cancer. Nevertheless, the primary types of cancer-fighting treatments, such as surgery, chemotherapy and radiation remain the usual options. However, besides their typical side effects and hazards, due to the high chance of relapse and TNBC’s partial response to the above-mentioned approaches, alternative therapeutic routes are urgently required. In this context, a wide range of non-invasive and spatiotemporally controlled therapies based on various physical stimuli, such as magnetic fields and light [3], have been developed over the past years as solutions to more effectively fight TNBC. Compared with traditional therapeutic strategies, the pN-level magneto-mechanical force (MMF) generated by magnetic nanomaterials under the effect of an external low-frequency rotating magnetic field (RMF) can be regarded as a powerful tool for the local therapy of deep-seated tumours, their efficiency being augmented by important advantages such as safety, non-invasiveness, facile spatiotemporal manipulation and deep tissue penetration [4–6]. Magnetic nanoparticles (MNPs) exposed to such low-frequency RMF (0~100 Hz) can perform an intricate series of motions [7], including vibration and rotation, which generate mechanical forces sufficient to activate mechanotransduction pathways [8–10], such as changing the cytoskeleton morphology [11], regulating cell adhesion [12], and even inducing cell apoptosis or death [13–15]. In our previous work, we found that the response of magnetic ferrite nanoparticles to external magnetic fields can be significantly improved by the controlled doping of zinc ions [16]. Subsequently, we showed that such MNPs can be used to perform dual-modality therapy with magneto-mechanical force and chemotherapy can be jointly exploited to overcome the drug resistance of cancer cells [17].

Considering the severe toxic effects of chemotherapy on normal body tissues and its limited specificity for cancer cells [18], photothermal therapy (PTT), in which photothermal agents usually stimulated by near-infrared (NIR) light produce local heat in tumour tissue and eradicate pathological cells, with superior tissue penetration and selective destruction of tumour cells, is a more prospectively mini-invasive therapeutic approach [19,20]. Most PTT studies to date have focused on the use of light in the NIR-I region (750-1000 nm), while the NIR-II light (1000-1700 nm) window is emerging as a more desirable light source for PTT due to lower photon scattering in tissues, accounting for deeper penetration and more reduced phototoxicity for healthy tissues [21]. Additionally, in order to avoid harsh thermal treatment unfavourably activating the undesirable cell death process known as necrosis, which may trigger inflammatory reactions and cancer metastasis [22], mild NIR-II hypothermia (41-48 °C) allows cancer cells to be preferentially eliminated by apoptosis without damaging normal tissues, a feature that is highly important for clinical application [23,24]. Notably, the temperature modulation of hypothermia is intimately correlated with the expression of heat shock proteins (HSP) [25].

In this work, we demonstrate in both *in vitro* and *in vivo* settings that the combined use of these two complementary and synergistic therapeutic strategies, namely MMF and NIR-II hypothermia, represents an exquisite route for fighting TNBC. To this end, we have designed and developed a multifunctional magnetic-photothermal converter (MPC) consisting of zinc-doped ferrite nanoparticles (Zn_0.2_Fe_2.8_O_4_ NPs) and polyethene glycol (PEG). We demonstrate that the MMF generated by the highly efficient magnetic field-mediated MPCs can dramatically inhibit the migration and invasion of MDA-MB-231 cells, and that this therapeutic route can be applied in tandem with NIR-II hypothermia, to result thus in a “double effect” strategy. Furthermore, we show that these two therapeutic layers are synergistic, MMF augmenting the outputs of NIR-II hypothermia. To shed light on this, we comprehensively investigate the mechanisms of the MPCs-mediated magneto-photothermal synergistic therapy (MPST) to clarify the intrinsic link between the two treatment modalities (**Scheme 1**). Given the excellent results of this “double effect” therapeutic scheme and the considerable biocompatibility and biosafety profiles of the MPCs, we consider that the proposed framework holds important potential to pave the way for some future cutting-edge clinical approaches that can successfully fight TNBC as well as cancers in general.

**Scheme 1.**
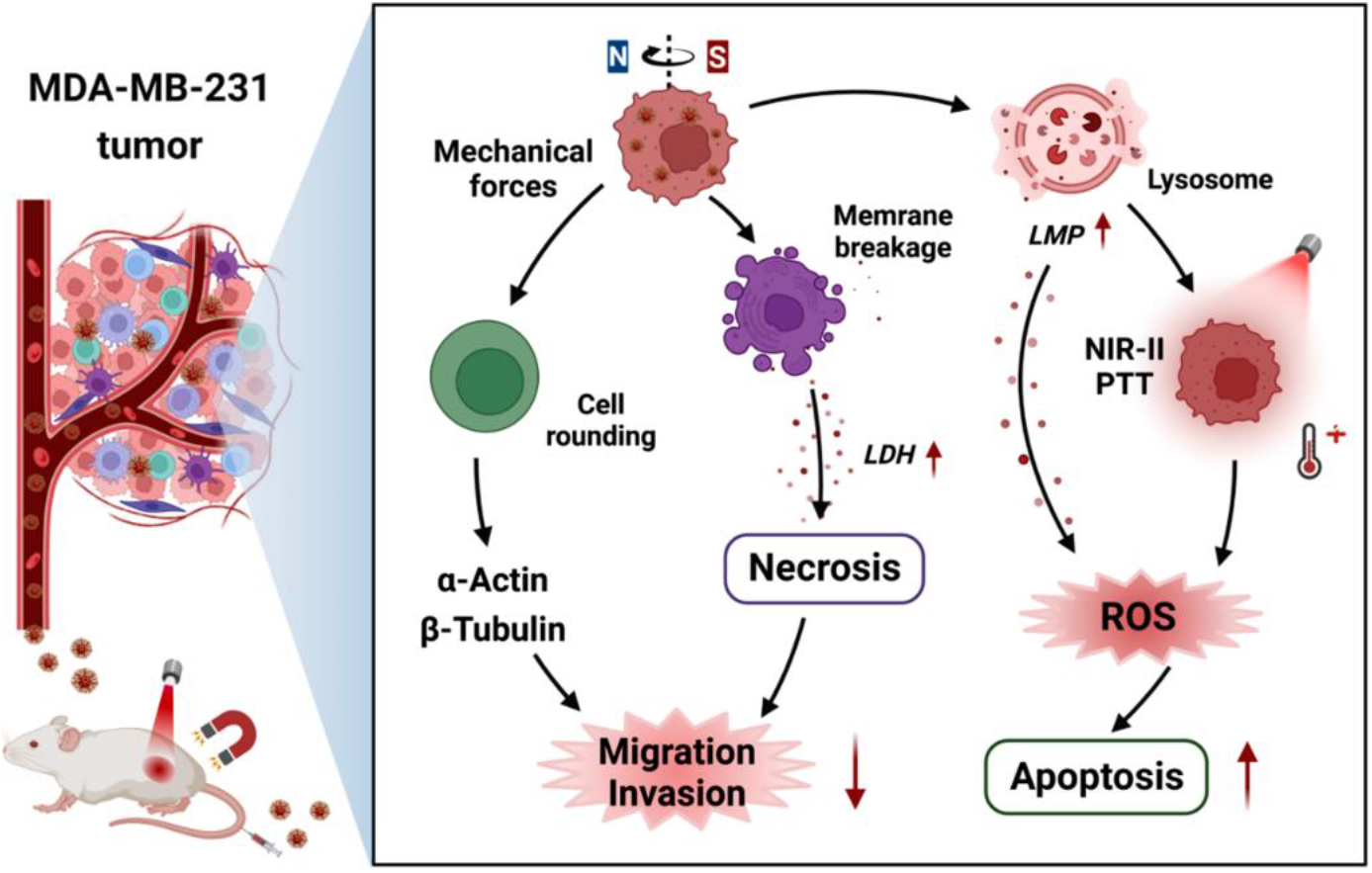
Schematic illustration of the MPCs-mediated magneto-photothermal synergistic strategy (MPST) for TNBC therapy. The zinc-dopped ferrite nanoparticles were modified to obtain biocompatible nanocomposites (MPCs) with excellent magnetism and NIR-II photothermal abilities. After being internalized into TNBC cells, MPST treatment was carried out. Under a 15 Hz rotating magnetic field, the generation of MMF was found to effectively inhibit the migration and invasion of MDA-MB-231 cells, while also enhancing the lysosomal membrane permeability and elevating the intracellular ROS, making cells sensitive to the subsequent heating. The second therapeutic layer based on exposure to NIR-II hypothermia was found to elevate intracellular oxidative stress, accelerating cell death. The synergistic effect between MMF and NIR-II PTT was demonstrated as being highly effective both *in vitro* and *in vivo*.

## 2. Results and Discussion

### 2.1. Design and Characterization of the MPCs

The physicochemical properties of the MNPs determine their magnetic responsiveness and their NIR photothermal effect. To determine these, a rich palette of investigations has been performed. Approx. 20 nm hydrophilic zinc-dopped ferrite nanoparticles were obtained by the high-temperature hydrothermal method and modified with PEG to enhance their dispersibility and biocompatibility (Figure **1a**). Transmission electron microscopy (TEM) revealed that the MPCs exhibited comparatively good dispersion, with an average size of 80 nm (Figure **1b** and Figure **S1**). It should be noted that larger magnetic particles have the potential to generate a more powerful magnetic force, and consequently yield more efficient therapeutic effects compared to smaller ones. High-resolution transmission electron microscopy (HR-TEM) and fourier transform infrared spectroscopy (FTIR) demonstrated the successful modification of PEG molecules on the surface of the MPCs (Figures **1c**, **g**). The selected area electron diffraction (SAED) mode image showed different diffraction rings, indicating a highly crystalline structure (Figure **1d**). As shown in Figure **1e** and Table **S1**, according to elemental mapping analysis (EDS) and inductively coupled plasma optical spectroscopy (ICP-OES), the ferrites in the MPCs are Zn_0.2_Fe_2.8_O_4_ NPs. The X-ray diffraction (XRD) patterns in Figure **1f** confirmed the spinel cubic structure of the MPCs (JCPDS card no. 22-1012). In addition, the saturation magnetization (Ms) results showed that the MPCs exhibit weak ferromagnetic behaviour with a coercivity of 32 Oe, while the Zn doping yields an optimal Ms, reaching 86.7 emu/g (Figure **1h**) at room temperature, indicating that MPCs can display superb responsiveness to an applied rotating magnetic field.

**Fig. 1.**
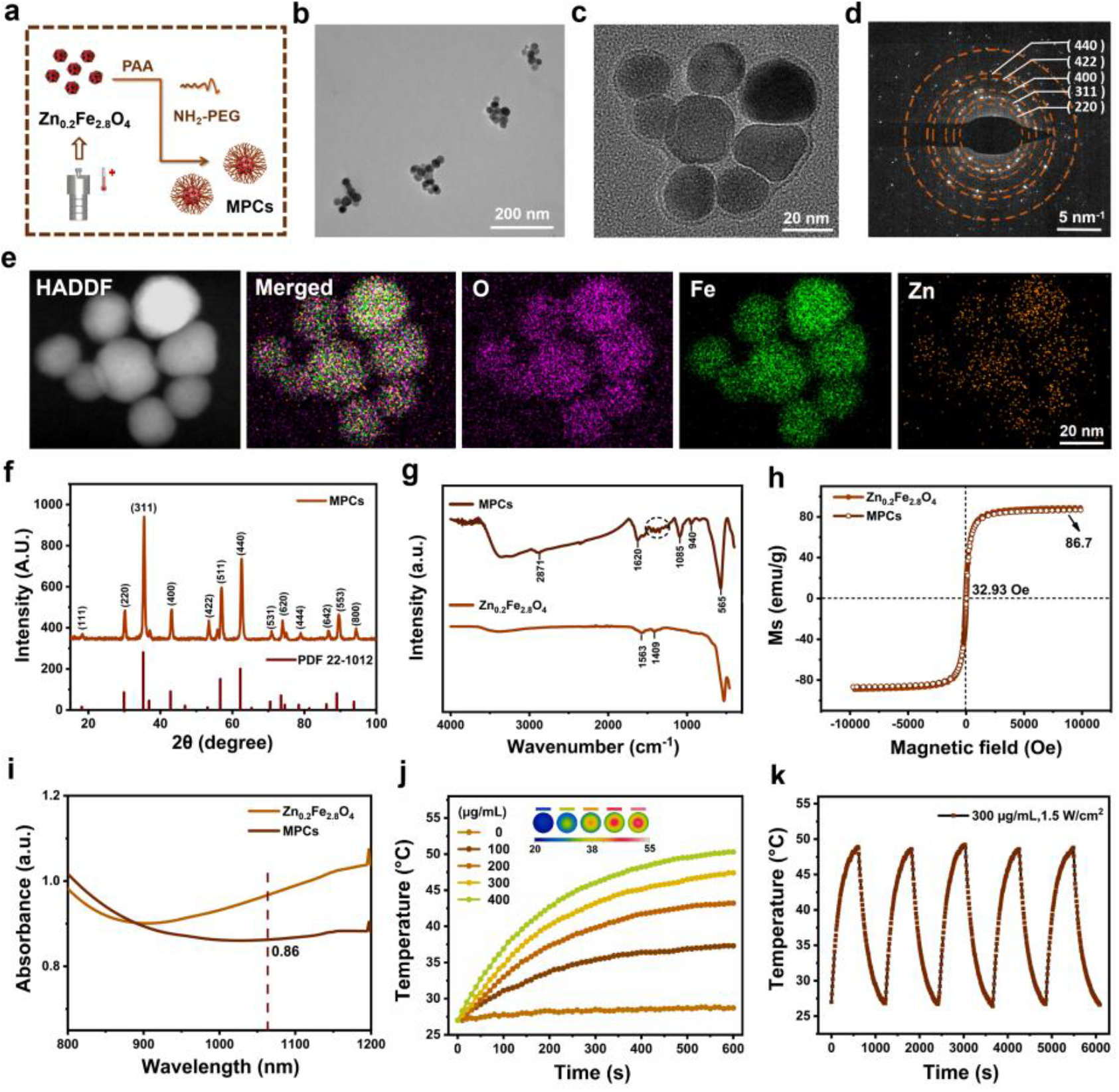
Synthesis and characterization of the MPCs. (a) Schematic illustration of the preparation process. (b) TEM image of MPCs. (c) HR-TEM image of MPCs showing that the surface of MNPs is wrapped with a PEG layer of uniform thickness. (d) The SAED mode. (e) EDS analysis showing that the Zn, Fe and O species were uniformly distributed on the individual MPC nanoparticles. (f) XRD spectrum at room temperature. (g) FTIR spectra demonstrating the successful modification of PEG on the surface of Zn_0.2_Fe_2.8_O_4_ NPs, where 940 cm^−1^ and 1085 cm^−1^ both represent the bending vibration peaks of the C-H bond, 1620 cm-1 and 2871 cm-1 are indicated as the stretching vibration peaks of C=O and C-H bonds, respectively, and the dashed circle indicates the C-C skeleton vibration peaks. (h) Ms results and (i) NIR absorption spectra of MPCs (300 μg/mL) and Zn_0.2_Fe_2.8_O_4_ NPs (300 μg/mL) at 300 K, respectively. (j) Photothermal conversion of MPCs at 1064 nm (1.5 W/cm^2^) at different concentrations (0, 100, 200, 300, 400 μg/mL). (k) Temperature variation over 5 on/off cycles of laser excitation.

An additional set of investigations showed that the MPCs exhibit significant absorption in the NIR-II window (Figure **1i**). Under 1064 nm laser light irradiation (1.5 W/cm^2^) for 10 min, the aqueous solution of MPCs showed a dramatic increase in temperature of 20 °C (Figure **S2a**). Further assays indicated that the temperature rise profiles and infrared images of MPCs were positively correlated with the concentration of MPCs and laser power density, revealing their controllable photothermal behaviour (Figure **1j** and Figure **S2b**). Remarkably, the MPCs exhibited excellent thermal stability after five heating and cooling cycles under laser irradiation (Figure **1k**). According to the change in temperature, the photothermal conversion efficiency (PCE) in the NIR-II region was calculated at 11.65% (Figure **S2c**).

### 2.2. MMF effects on Cell Fate

To objectively assess the extent of cancer cell damage achieved by MMF effects generated by the MPCs under an RMF, the intrinsic cytotoxicity of MPCs was first investigated. Over 80% of MDA-MB-231 cells behaved bioactive when incubated with MPCs (800 μg/mL) for 24 h (**Figure 2a**). As shown in **Figure 2e**, MPCs (300 μg/mL) were completely internalized by MDA-MB-231 cells after 12 h of co-incubation. TEM imaging showed that the MPCs mainly tend to localize in organelles such as lysosomes and partially in the cytoplasm (Figure **2f**). Under the demonstrated premises of excellent biocompatibility, the MMF effects of the MPCs were further investigated *in vitro*.

**Fig. 2.**
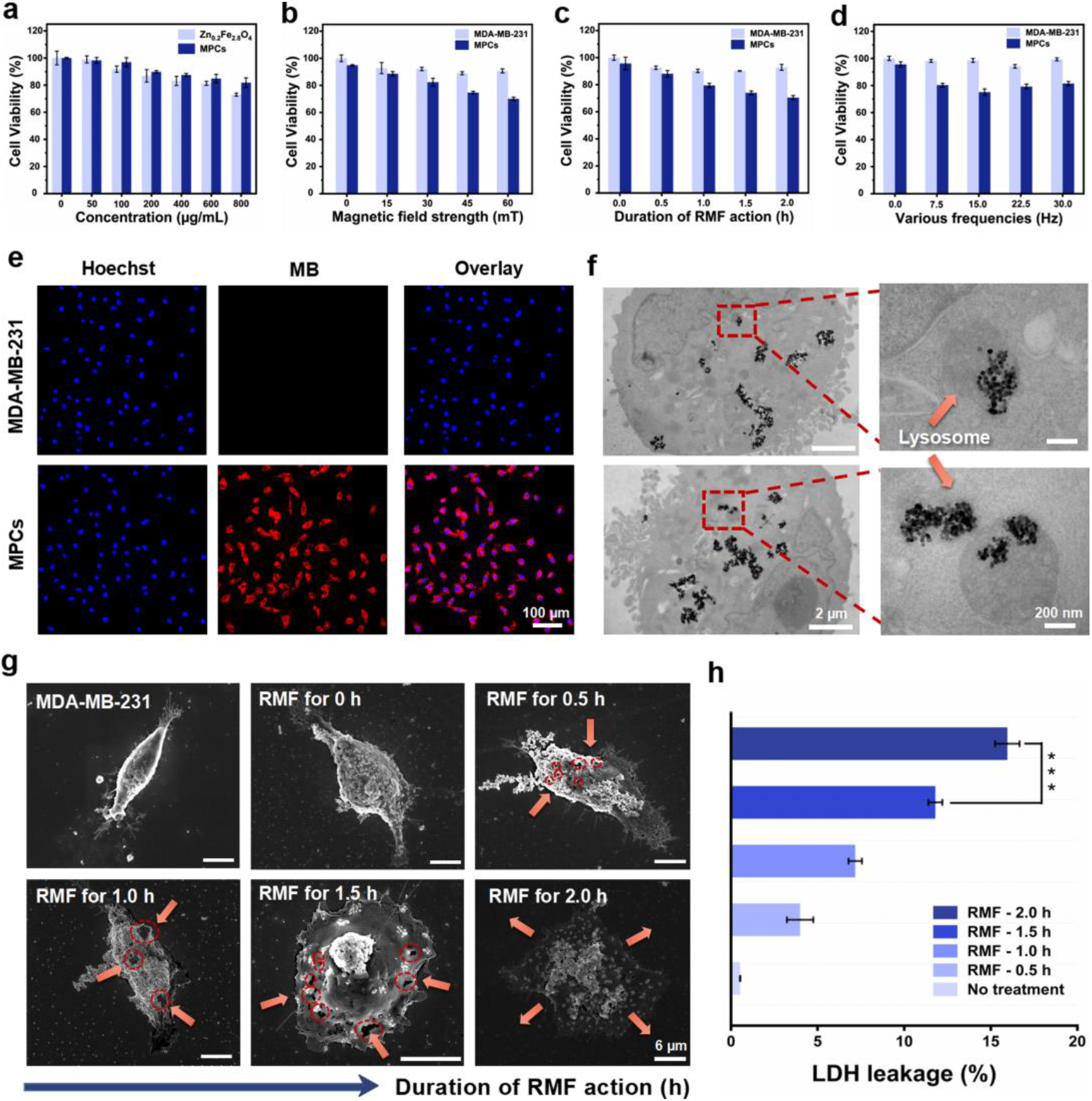
MDA-MB-231 cancer cell death induced by MPCs by magneto-mechanical force effect. (a) Cell viability of the MDA-MB-231 after co-incubation with different concentrations of MPCs for 24 h. Cell viability under the RMF of different strengths (b), durations (c), and frequencies (d) of RMF, respectively. (e) The uptake of MPCs (300 μg/mL) by MDA-MB-231 cells following 12 h of co-incubation (Scale bar: 100 μm). (f) TEM images showing the localization of MPCs within MDA-MB-231 cells (Scale bar: 2 μm; Zoom in: 200 nm). (g) SEM images of cells under various RMF exposure intervals (Scale bar: 6 μm). (h) The release of LDH from cells after treatment with MPCs (300 μg/mL) upon exposure to RMF with different action time. \*\*\**P* < 0.001 indicates a statistical difference between the two groups. Data are expressed as mean ± standard deviation. Error bars are based on the results of at least three independent experiments per group.

As shown in Figures **2b**, **c**, the strength and the action time of the magnetic field were found to play important roles in destroying cancer cells under the same treatment conditions, indicating that the stronger the magnetic field or the longer the action time of the RMF, the lower the cell viability. Conversely, exposure of pure cells (without internalized MPCs) to RMF did not inhibit their viability (Figures **2b, c**). Interestingly, under the effect of RMF at 60 mT for 2 h, the cell viability showed a trend of decreasing at first and afterwards increasing with the increased magnetic field frequency in Figure **2d**, indicating that MPCs and RMF reached the maximum synchronization frequency of 15 Hz [26]. Hence, the mechanical killing of MPCs under this setting displayed the strongest mechanical performance with an optimal cell lethality ratio of 23%.

Furthermore, scanning electron microscopy (SEM) imaging was used to reveal how the extent of cell membrane damaged by MMF correlates with the RMF action time (Figure **2g**). The perforation of the cell surface became more obvious with increased RMF exposure time, as evidenced by the significantly increased number and diameter of the cellular membrane holes, followed by complete cleavage of the cell membrane in part of the cells after being treated with RMF for 2 h. Additionally, damage to the cell membrane was also found to cause the release of important intracellular components such as lactate dehydrogenase (LDH). In this respect, Figure **2h** shows that the release of intracellular LDH increases dramatically over the duration of RMF. Considering that the perforation of the cell membrane and the consequent discharge of more intracellular components eventually leads to cell death, the therapeutic effects of MMFs can clearly be noticed.

### 2.3. MMF Effects on Cell Migration and Invasion

In comparison to assessing the potential of MPCs to kill cancer cells by MMF effects, objective quantification of their capacity to suppress the migration and invasion of tumour cells (Figure **3**) was significantly more challenging, given the high metastatic potential of the MDA-MB-231 cells (TNBC cell line). Such cells are vigorously aggressive, resistant to chemotherapeutic agents such as dasatinib [27], and exhibit unanchored growth independent of growth factors [28]. The cytoskeleton provides stability for cell morphology and participates in critical processes such as cell mitosis, endocytosis and motility, performing an essential role in the migration and invasion of tumour cells, for which actin filaments and microtubules are the main components [29]. Based on this, as shown in Figures **3a**, **b**, β-tubulin and F-actin were stained by immunofluorescence and ghost pen loop peptide, respectively. Using confocal laser scanning microscopy (CLSM), untreated MDA-MB-231 cells, with their characteristic shuttle-shaped growth pattern, were found to be structured by organized β-tubulin and F-actin architecture with some filopodia or lamellipodia formation (Figures **3a**, **b**). As the RMF (60 mT, 15 Hz) exposure time was prolonged, the cells treated with MPCs crumpled into round shapes, which was accompanied by remarkable changes in the cytoskeleton, resulting in more skeletal proteins being focused on the outer edges of the cells and the disappearance of the cell membrane folds, together with the filamentous and lamellar pseudopods.

**Fig. 3.**
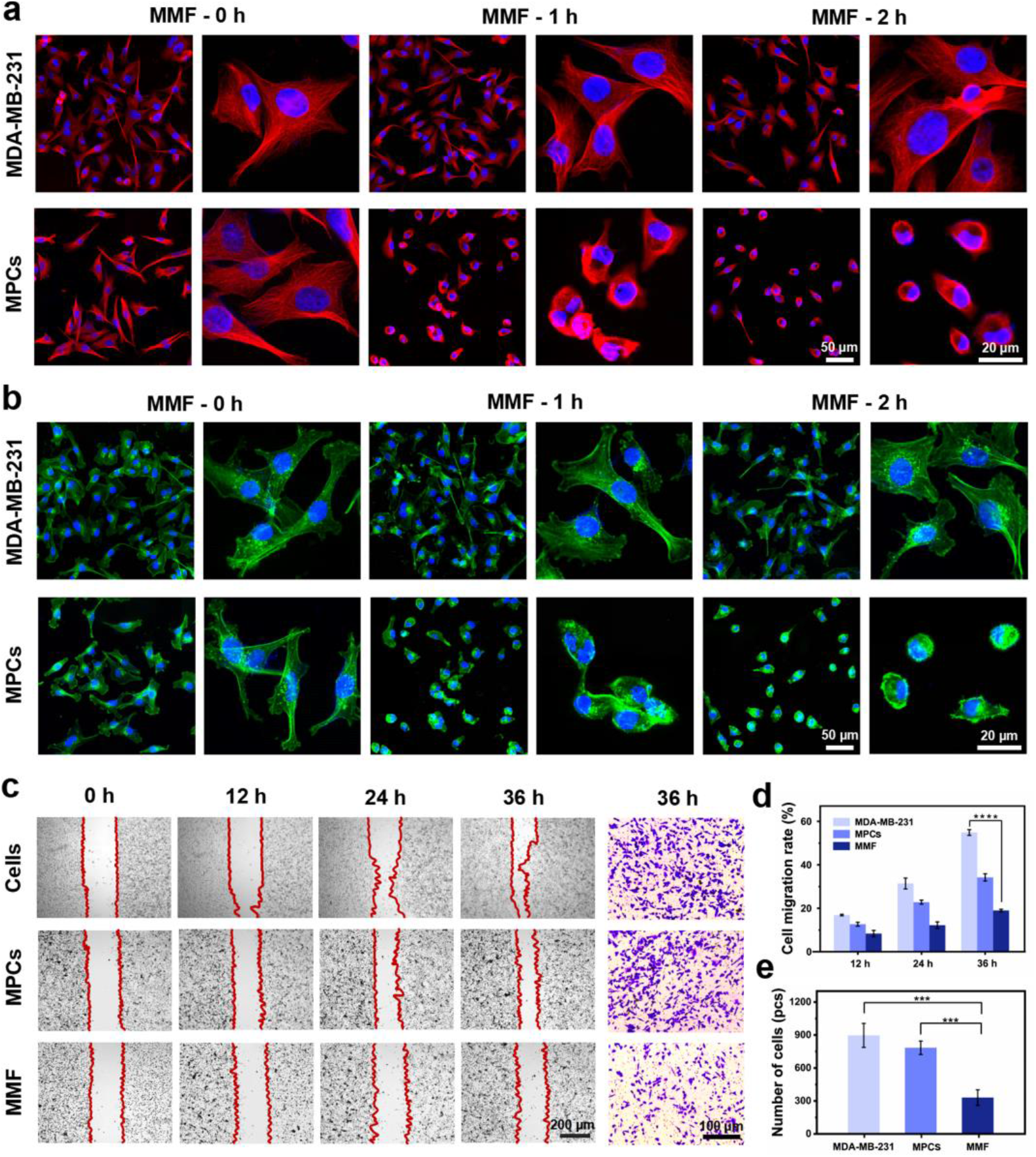
MMF effects generated by the MPCs under RMF modify the cytoskeleton in a manner that inhibits cell migration and invasion. Representative CLSM images of (a) microtubule network and (b) actin cytoskeleton in control and MPCs treated MBA-MD-231 cells at various RMF exposure time. β-tubulin (red), F-actin (green) and DAPI-stained nuclei (blue). Scale bar: 50 μm; Zoom in: 20 μm. (c) The cell wound healing and transwell invasion assays of MDA-MB-231 cells treated with MMF effects correspond to the MPCs under RMF (the black scale bar: 200 μm and 100 μm, respectively). (d) Cell wound healing assay: evaluation of migration distance within 36 h (n=3). (e) Transwell invasion assay: number of cells crossing the chambers over 36 h (n=3). \*\*\**P* < 0.001, \*\*\*\**P* < 0.0001 indicates a statistical difference between the two groups. Data are expressed as mean ± standard deviation. Images are representative of three independent experiments.

The significant changes that we observed in the cytoskeleton structure, motivated us to further evaluate the effect of MMF on the migration and invasion of MDA-MB-231 cells (Figure **3c**), which we accomplished by cell wound healing and transwell invasion assay. The migration ability of cancer cells was assessed by the average decrease in the distance between wound edges at different time points (0 h, 12 h, 24 h, 36 h) in the presence or absence of MMF stimulation. The cell wound healing assay results displayed in Figures **3c**, **d** and Table **S2** revealed that cells treated with MMF significantly repressed wound healing with a 67.63% decrease in cell migration compared to the pure cell group at 36 h. Consistent with this finding, we also identified by transwell invasion assays (Figures **3c**, **e** and Table **S3**) that the invasive capacity of the cells dramatically decreased to 71.82% of the original level within 36 h after MMF treatment. Our investigations showed that treatment of MDA-MB-231 with MPCs has no influence on cell migration and invasion in the absence of RMF.

### 2.4. MPST Effects on Cell Fate in Vitro

In previous sections, we showed that the MMF generated by the MPCs under RMF exposure inhibits the growth, migration and invasion of tumour cells to some extent. Here we demonstrate a strategy for using MMF in concert with NIR-II Hypothermal Ablation, a second, complementary therapeutic route that augments the outputs of the first therapeutic layer, resulting in a highly efficient “double effect” weapon against TNBC, the MPST. To this end, we exploit the excellent NIR-II photothermal properties of the MPCs, and use low-temperature PTT in the NIR-II window (1064 nm) to eliminate surviving MDA-MB-231 cells that were previously treated by MMF. We observe that the effects of PTT are facilitated by prior MMF ones, which makes the two therapeutics highly synergistic.

Under 1064 nm laser irradiation (1.5 W/cm^2^), the viability of cells treated with MPCs was observed to decrease substantially with increased exposure time (Figure **4a**), while pure cells were virtually unaffected by the NIR light irradiation, corresponding to the infrared images shown in Figure **4b**. To further evaluate the effectiveness of the MPST strategy, the MPST group was conducted at different PTT times (0, 4, 8, 12 and 16 min) with an optimized MMF (15 Hz, 60 mT, 2 h), where the calculated Q values exhibited an increase then decrease trend, reaching a maximum of 1.39 at an irradiation time of 8 min, which is higher than the determined threshold for the synergistic effect index of 1.15 [14] (Figure **4c** and Table **S4**). Therefore, the treatment time in the subsequent *in-vitro* experiments was set to the optimized value of 8 min for PTT, accompanied by a temperature increase to 47 °C, at which the enhancement of intracellular heat resistance due to HSP overexpression could be effectively avoided (Figure **4b**) [30,31]. The CCK-8 assay results displayed in Figure **4d** showed that over 80% of cancer cells were killed after MPST treatment, which had a considerably higher killing rate than those observed for conventional MMF (23%) and PTT (40%). These results reflect the valuable potential of the synergies occurring between MMF and PTT therapies.

**Fig. 4.**
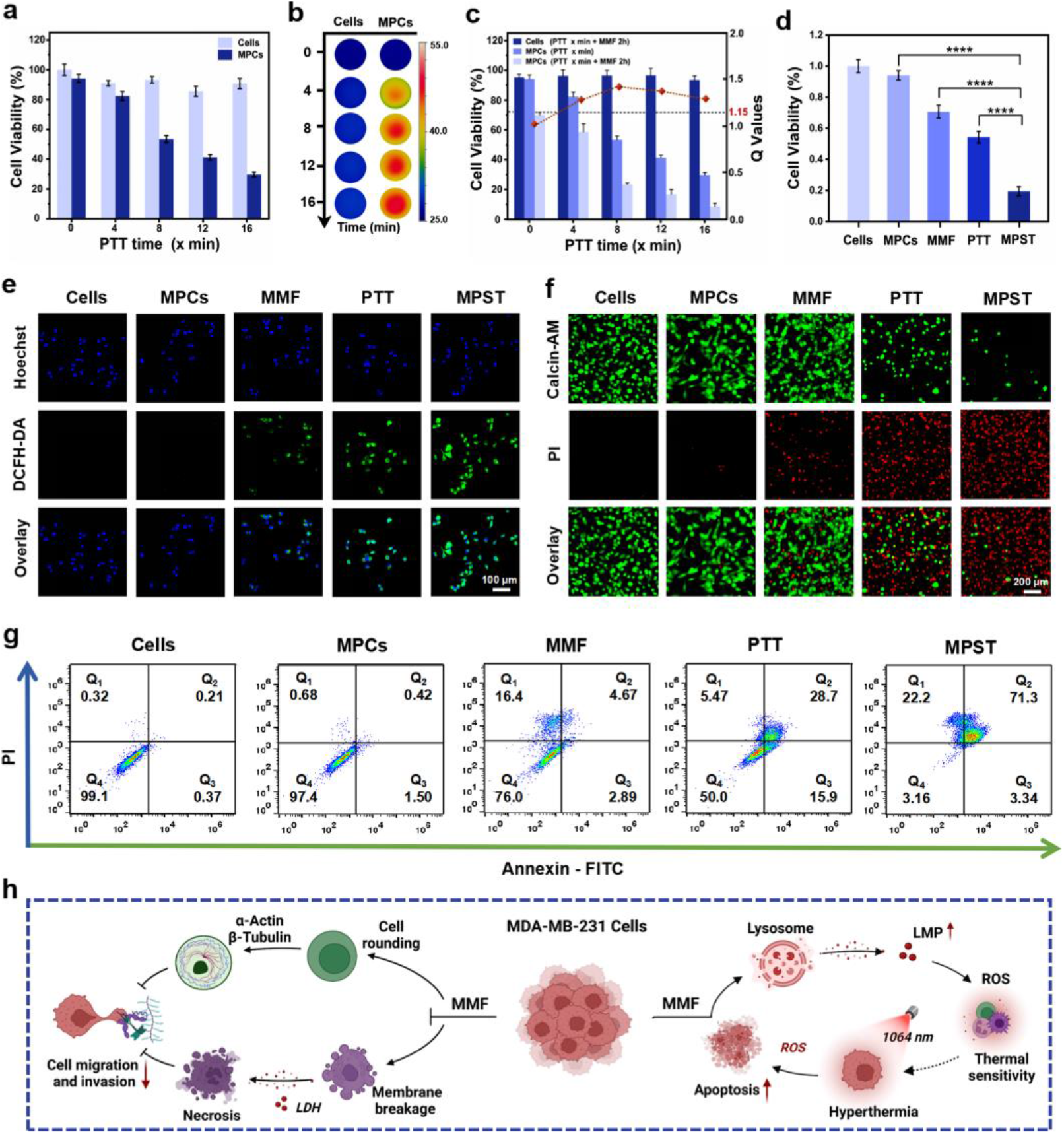
MPST effect: MPCs induce MDA-MB-231 cell death by in-tandem, synergistic, use of MMF and NIR-II hypothermia. *In vitro* viability (a) and infrared images (b) of cancer cells treated with MPCs-enabled (300 μg/mL) NIR-II hypothermia at different irradiation times (0, 4, 8, 12, 16 min) with a 1064 nm laser. (c) The viability of cancer cells and Q values under MPST treatment consisted of 2 h MMF and varied PTT time. Q ≥ 1.15 demonstrate the existence of a synergistic effect between the two therapeutic routes. (d) Therapeutic effects on MDA-MB-231 cells of MPCs-based MMF, PTT and MPST therapies assessed by the Cell Counting Kit assay. Relevant treatment parameters are RMF with 15 Hz, 60 mT, 2 h and PTT with 1.5 W/cm^2^, 8 min. Statistical analysis was obtained using a t-test, \*\*\*\**p*<0.0001 (n ≥ 3). (e) CLSM images of ROS levels in MDA-MB-231 cells treated with MPCs upon different therapeutic modalities (Scale bar: 100 μm). (f) The Calcein AM/PI-costained of cells after various treatments (Scale bar: 200 μm). (g) The AnnexinV-FITC/PI analysis of cell death patterns under exposure to diverse treatments using flow cytometry. (h) Mechanistic diagram of the MPST therapy *in vitro*.

To further investigate the mechanism by which the “double effect” MPST therapeutic strategy achieves cancer cell death, CLSM imaging was employed to evaluate the overall level of reactive oxygen species (ROS) of *in vitro* MDA-MB-231 cells stained with a 2,7-dichlorodi-hydrofluorescein diacetate (DCFH-DA) probe (Figure **4e**). Besides the significantly elevated ROS levels of the PTT and MPST treated cells, we also noticed an unexpected increase in the intracellular ROS levels after MMF treatment, suggesting that the mechanical stimulation of cells can effectively up-regulate their biochemical ROS signalling, which we hypothesized that may be correlated with an enhanced permeability of the lysosomes [32,33]. To validate our hypothesis, the changes in the subcellular structure and lysosomal permeability (LMP) were assessed using acridine orange (AO) fluorescence detection reagents. Generally, in intact lysosomes, AO exists as a protonated oligomer with red fluorescence: the manifested red fluorescence spots gradually diminish with the increase of LMP (Supplementary, Figure **S3**). In comparison with the pure cell group (without internalised MPCs) and the MPCs group without exposure to RMF or NIR light, a significant weakening of the red fluorescent spots was observed in the MMF and PTT groups, implying that individual MMF or PTT treatments were able to disrupt the lysosomal membrane by mechanical force or heat, respectively. Furthermore, for the MPST group, the red fluorescent spots were no longer visible, indicating a greater extent of LMP.

The aggravated oxidative damage to tumour cells in the mid-to-late stage of apoptosis and death under various treatments was further verified by Calcein Acetoxymethyl Ester (AM)/propidium iodide (PI) costaining (Figure **4f**). To further explore the mechanism of cancer cell death after treatments with MMF, PTT and MPST, we used Annexin V-fluorescein isothiocyanate (FITC)/PI costaining by flow cytometry (Figure **4g**), where AnnexinV can conjugate to phosphatidylcholine (PS) and translocate from the interior to the exterior of the plasma membrane during early apoptosis, while PI is utilized to assess the integrity of the cell membrane, signalling late apoptotic or necrotic cell death [34,35]. According to these analyses, the cell death mode of the MMF group was primarily mechanical necrosis, while for the PTT and MPST groups, late-stage apoptosis was predominant.

From the available results, it can be concluded that in the presence of RMF, the MMFs generated by the massive accumulation of intracellular MPCs cause direct mechanical destruction to the cell membrane, leading to partial cell mechanical necrosis and further inhibiting cell migration and invasion. The MMF effects were also shown to disrupt lysosomes, causing a massive release of internal proteolytic enzymes and acids, which lowers the cytoplasmic pH and induces oxidative stress reactions in cells, thereby making them more sensitive to external stimuli (*e.g*., heat). Thus, the therapeutic potential of subsequent hypothermia in the NIR-II region was considerably augmented, accompanied by high levels of ROS generation, hence accelerating the apoptosis of cancer cells (Figure **4h**).

### 2.5. Therapeutic Efficiency of MPST in Vivo

Following the exploration of the MPST mechanisms occurring at the cellular level *in vitro*, an MDA-MB-231 subcutaneous tumour model was constructed and applied to evaluate the therapeutic effects of MPST *in vivo*. When the tumour volume reached 100 mm^3^, the MPCs were administered intravenously to mice three times on days 0, 3 and 6 with a dose of 30 mg/kg per injection, satisfying the biosafety profile (Figure **5a**, Figures **S4**, **5**). The accumulation of MPCs at the tumour site was indirectly determined by monitoring its heating upon 1064 nm laser irradiation every 2 h. After 8 h of MPCs injection, the temperature rose significantly, and following this time point, no additional increase could be observed, which demonstrated that the accumulation of MPCs at the tumour site reached the maximum level at this point (Figure **S6**). Accordingly, after 8 h of intravenous injection, the MPCs-mediated treatments were performed three times every two days, from day 0 to day 6 with treatment parameters of 60 mT, 15 Hz, and 2 h for MMF and 1.5 W/cm^2^, 15 min for PTT, respectively. Noteworthy, the RMF parameters and NIR-II laser power in this study were both positioned in the safe range [36,37].

**Figure 5.**
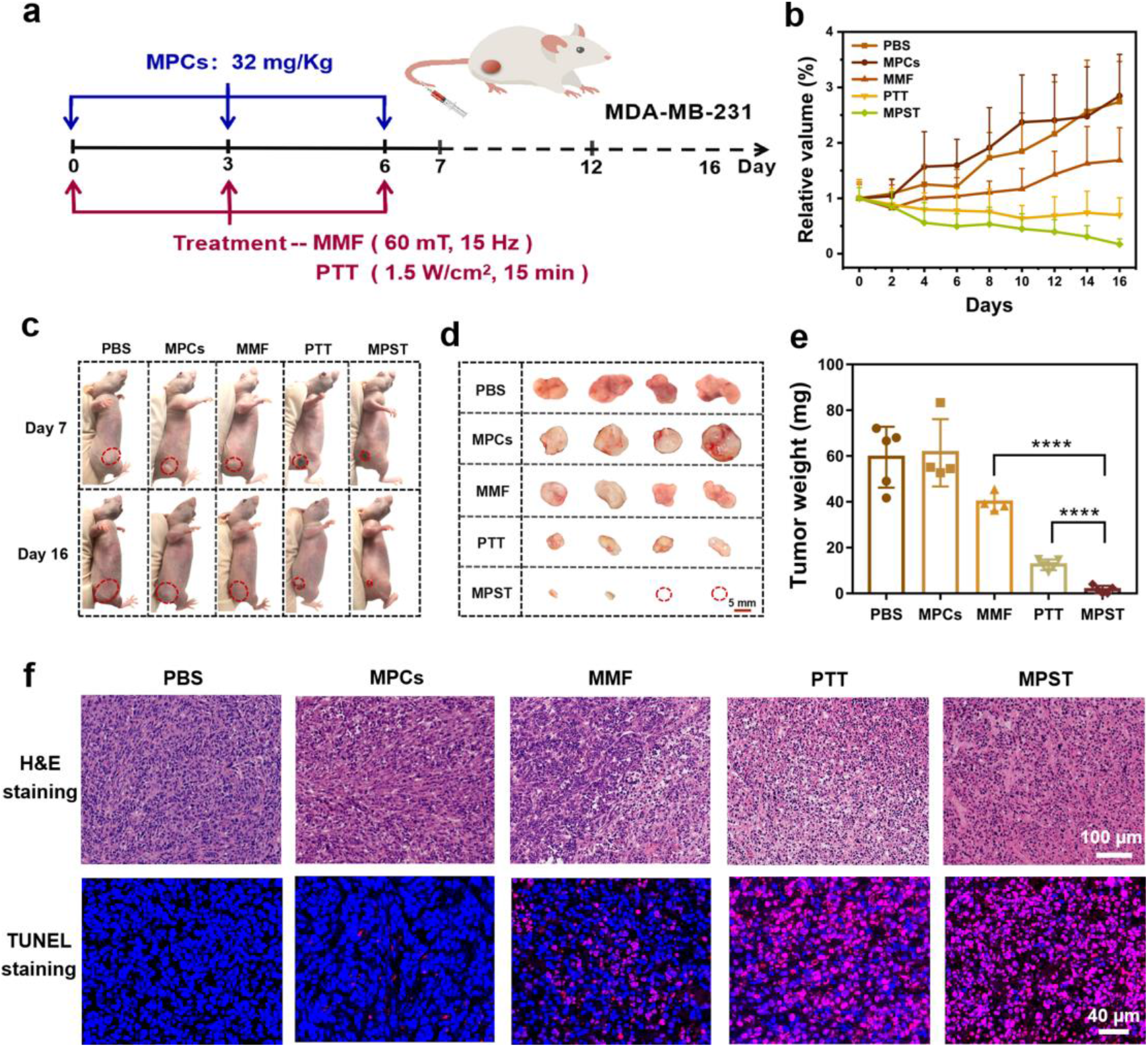
The therapeutic effectiveness of MPST in MBA-MD-231-bearing mice. (a) Schematic diagram of the treatment procedure *in vivo*. MPCs were injected intravenously and treated on days 0, 3 and 6 for three times, with the parameters of 60 mT, 15 Hz, and 2 h for MMF and 1.5 W/cm^2^, 8 min for PTT. (b) Relative tumour volume of mice in various treated groups (n ≥ 5). (c) The pictures of mice with tumours on day 7 and day 16. Pictures (d) and weights (e) of tumours from each treatment group were obtained after dissection (Scale bar: 5 mm). Statistical analysis was obtained using a t-test, \*\*\*\**p* < 0.0001 (n ≥ 4). (f) H&E staining and TUNEL staining of pathological changes in tumour tissues from various treatments, where TUNEL staining can detect apoptosis of cells. FITC-dUTP stained cells in apoptosis (red) and DAPI-stained nuclei (blue). Scale bar: 100 μm and 40 μm.

The body weight and tumour size of MDA-MB-231-bearing mice were monitored daily for 16 days. The body weight of the mice was maintained essentially steady (Figure **S7**, supporting information). MMF had an inhibitory effect, while PTT was enhanced and significantly reduced the tumour size. Importantly, tumours in the MPST group were entirely eliminated (Figures **5b-d**). Moreover, the quantitative analysis of tumour weight was consistent with the tumour images (Figure **5e**). Additionally, Hematoxylin-Eosin (H&E) and TdT-mediated dUTP nick end labelling (TUNEL) staining revealed that the tumour tissue was found to be entirely necrotic under MPST treatment, MMF alone did not prevent cancer cells from developing into tumours whereas low-temperature PTT remarkably reduced tumour size, accompanied by massive tumour tissue necrosis (Figure **5f**).

To investigate the metabolism of MPCs after 16 days in mice, Prussian blue dye was applied to stain the MPCs. Fluorescent signals corresponding to this contrast agent were not observed at the tumour site, while signals originating from the liver and spleen indicated some extent of MPC accumulation in these organs (Figure **S8**). However, the overall results obtained by H&E staining, blood routine and blood biochemical analysis, displayed in Figures **S9**, **10**, no apparent MPCs-induced damage was discovered in the heart, liver, spleen, lung, and kidney tissues, indicating the safety of this MPST enabling therapeutic agent. The *in vivo* results reveal thus that the MPCs-mediated MPST strategy has a significantly destructive effect on breast tumours and this treatment approach is likely to have minimal impact on the health state of the patient, according to biosafety assessment in mice, rendering this MPST with MPCs framework as a potential candidate for a future cutting-edge clinical therapy for fighting TNBC and cancers in general.

## 3. Conclusion

In summary, to efficiently treat TNBC, we designed and demonstrated a custom-tailored strategy exploiting the synergisms occurring between therapeutic effects of magneto-mechanical forces and photo-thermal damage induced by MPCs, featuring the superb capacity to accurately respond to external physical stimuli. We have first developed a versatile MPC with excellent biocompatibility as a force-thermal converter, capable of transforming external magnetic and light in the NIR-II window into force and heat in a well-controlled manner. Second, we demonstrated that the MMF generated by MPCs under the effect of RMF significantly diminished the migration and invasion capacities of tumour cells by 67.63% and 71.82%, respectively, *via* changing the cytoskeleton morphology while exerting a significant inhibitory effect on the growth of MDA-MB-231 cells. Furthermore, we also found that the MMF generated by the massive accumulation of intracellular MPCs caused direct mechanical destruction to the cell membrane, leading to partial cell mechanical necrosis, and disruption of intracellular lysosomes. This latter outcome resulted in increased LMP levels and the related release of large amounts of protein hydrolases, which further induced intracellular oxidative stress. These results were found to be synergistic with the NIR-II hypothermal effects, enhancing their extent and significantly augmenting the outputs of the PTT strategy. Applying the PTT treatment after MMF led to significantly more elevated levels of intracellular ROS compared to the case where MMF was not previously applied, and was found to massively accelerate the induction of apoptosis. Therefore, the MPST strategy leads to a chain reaction of the cell membrane, skeletal proteins, subcellular organelles, and biological signals, which results in remarkable therapeutic success against TNBC in both *in vitro* and *in vivo* settings. Although tested on TNBC, the proposed framework has high potential to be generalized to successfully fight against other cancer variants, in a less invasive and hazardous way than current state-of-the-art methods.

## 4. Materials and Methods

### 4.1. Materials and reagents

Ferrous sulfate heptahydrate (FeSO_4_·7H_2_O), sodium citrate, 1-(3-Dimethylaminopropyl)-3-ethylcarbodiimide hydrochloride (EDC) and N-hydroxy succinimide (NHS) were purchased from Aladdin Industrial Corporation (Shanghai, China). The zinc acetate dihydrate (Zn (Ac)_2_·2H_2_O) was obtained from Energy Chemical (Shanghai, China). Monohydrazine (N_2_H_4_·H_2_O) was purchased from Sinopharm Chemical Reagent Co., Ltd. (Shanghai, China). Polyacrylic acid (PAA, MW: 2000 Da) and Methylene blue (MB) were obtained from Macklin (Shanghai, China). Amino-modified polyethene glycol (mPEG-NH_2_, MW: 2000 Da) was purchased from Yare Biotech, Inc. (Shanghai, China). Cell Counting Kit-8 (CCK-8) was purchased from Apex BIO (America). Hoechst 33342 was obtained from Biosharp (Anhui, China). Reactive Oxygen Species Assay Kit (SOO33S), Calcein/PI Cell Viability/Cytotoxicity Assay Kit (C2015S), Lactate dehydrogenase cytotoxicity (LDH) assay kit (C0017) and Annexin V-FITC Apoptosis Detection Kit (C1067S) were obtained from Beyotime (Shanghai, China). The lysosomal membrane permeability (LMP) assay kit was obtained from Chen Gong Biotechnology Co., Ltd. (Shanghai, China). All other chemicals and solvents are of analytical grade (AR) and can be used directly without further purification.

### 4.2. Synthesis of the Zn_0.2_Fe_2.8_O_4_ MNPs

First, 1.8 mmol FeSO_4_·7H_2_O and 3.64 mmol Zn (Ac)_2_·2H_2_O were dissolved in 20 mL of deionized water, and 1.94 mmol citric acid was added quickly *via* magnetic stirring to obtain a uniform solution. At this point, 2.5 mL of 10 mol·L^−1^ N_2_H_4_·H_2_O was added dropwise to the mixture and mechanically stirred for 30 min. Then, the mixture was transferred to a 50 mL capacity Teflon-lined autoclave and heated at 200 °C for 16 h *via* using a high-temperature oven and naturally cooled to room temperature. In a further step, the supernatant was removed, and the precipitate was centrifuged at 8,000 rpm for 7 min and washed 6 times with alternating distilled water and absolute ethanol. Finally, it was stored by freeze-drying for 12 h and then protected from light.

### 4.3. Synthesis of the MPCs

Initially, 5 mL of the prepared Zn_0.2_Fe_2.8_O_4_ MNPs (2 mg/mL) were dispersed into 10 mL of PAA (5 mg/mL) under sonication conditions and mechanically stirred for 2 h. Secondly, the reactants were centrifuged and washed three times with deionized water. After that, 50 mg of mPEG-NH2 was added to it and sonicated for 30 min. Then, 15 mg each of EDC and NHS were added sequentially and sonicated for 30 min. Finally, after 12 h of mechanical stirring, the reaction products were washed three times with deionized water to obtain MPCs.

### 4.4. Cellular Viability Using the MMF Assay

The CCK-8 assay was used to assess the effect of treatment *via* MMF. Briefly, cells were seeded on 96-well plates (1 × 10^4^ cells per well) and incubated for 12 h to adhere to the well. Then, MPCs (300 μg/mL) were added and co-incubated for 12 h and treated using a magnetic rotation platform with different magnetic field strengths (0, 15, 30, 45, 60 mT), different action times (0, 0.5, 1, 1.5, 2 h) and different rotation frequencies (0, 7.5, 15, 22.5, 30 Hz). After this, the cells were put back into the incubator and continued to incubate for 4 h. CCK-8 (10 μL) was added and continued to incubate at 37 °C for 1.5 h. The absorbance at 450 nm was finally measured using an enzyme marker.

### 4.5. Analysis of Cell Morphology by SEM

The changes in morphological characteristics of MDA-MB-231 cells before and after magnetic treatment were analyzed by scanning electron microscopy. Briefly, MDA-MB-231 cells (1 × 10^5^ pcs/mL) were seeded on in silico (~ 1 × 1 cm^2^) and cocultured with MPCs (300 μg/mL) for 12 h. Secondly, redundant material was washed off and cells were fixed with 2.5% glutaraldehyde at 4 °C for 2 h, followed by 2 washes with PBS (10 min each). Cells were dehydrated with a gradient sequence of ethanol (30%, 50%, 70%, 90% and 100%) (15 min for each concentration). Finally, the prepared dried samples were observed by SEM at 10 kV.

### 4.6. LDH Release Assays

The results of LDH release from MDA-MB-231 cells cultured with MPCs were evaluated for 12 h at various times of RMF action. The absorption intensity at 490 nm of the cell culture medium containing the LDH test working solution was measured and the LDH leakage was calculated from LDH leakage (%) = [(absorbance of treated cells - absorbance of control cells)/ (absorbance of maximally enzymatically active cells - absorbance of control cells)] ×100. The assay was performed according to the corresponding instructions provided by the kit manufacturer.

### 4.7. Immunofluorescence Staining

To visualize the effect of the magnetic force on the cytoskeleton, immunofluorescence microscopy experiments were conducted on the MDA-MB-231 cell lines. Cells (2 × 10^5^ pcs) were seeded on crawl sheets (2 × 2 cm^2^) and co-incubated with MPCs (300 μg/mL) for 12 h. Secondly, they were treated with RMF of different action times and immediately fixed with 4% FBS for 30 min. Next, the slices were shaken dry and circled in the middle of the coverslip with a histochemical pen, and 50-100 μL of the film-breaking solution was added and incubated for 10 min at room temperature, washed 3 times with PBS for 5 min each time. Following this, cells were incubated with the primary antibody, anti-beta-tubulin mouse monoclonal antibody (1:200) at 4 °C overnight. After washing 3 more times with PBS, CY3-labeled secondary antibodies of the corresponding species were added to them and incubated for 50 min at room temperature, protected from light. Then, a TSA signal amplification reagent was added for 3-5 min, followed by 3 washes with PBS and 50-100 μL of FITC-labeled ghost cyclic peptide working solution was added to the cells and incubated for 2 h at room temperature. In addition, the nuclei of the cells were stained using DAPI. Finally, the slide was shaken dry with the cell side down and sealed on a slide with an anti-fluorescence quenching sealer, and placed under a fluorescence microscope (ECLIPSE E100, NIKON) for observation and image acquisition.

### 4.8. Cell Migration Assay

Firstly, MDA-MB-231 cells (1.2 × 10^6^ pcs) were seeded on a six-well plate containing a scratch assay insert. Next, the cells were co-incubated with MPCs (300 μg/mL) for 12 h, washed three times with PBS and replaced with serum-free DMEM. Afterwards, the scratch images were taken by a reverse fluorescence microscope (DMIL LED, Leica) at different time points (0, 12, 24 and 36 h) after magnetic treatment. Finally, the scratch area was calculated using Image J. The ability of MMF to inhibit cell migration (AMICM) is calculated by the following formula, where R_Cell_ and R_MMF_ represent the respective migration rates.

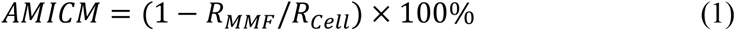

### 4.9. Cell Invasion Assay

The matrigel was first diluted to 250 μg/mL with serum-free DMEM at 1:40. Secondly, 100 μL of the diluted solution was applied to the upper layer of Transwell chambers and placed on a 24-well plate. After placing it in the incubator for 1 h to make the matrix gel sufficiently solidified, the upper chamber culture was aspirated and 200 μL of serum-free culture medium containing MDA-MB-231 cells (5 × 10^6^ pcs) was added to it. After that, the upper chamber was carefully placed with forceps on top of the lower chamber containing 15 % serum and placed back in the incubator for 36 h. After fixing with 4 % FBS at room temperature for 30 min, the upper chamber was washed twice with PBS and stained with 0.5% crystal violet for 15 min. The upper chamber was washed with PBS to remove excess crystal violet and wiped clean with a cotton swab. Finally, they were observed by light microscopy and counted by Image J. The ability of MMF to inhibit cell invasion (AMICI) is calculated by the following formula, where N_Cell_ and N_MMF_ represent the number of cell invasions, respectively.

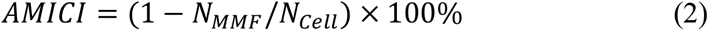

### 4.10. Effect of MPST

First, 1.2 × 10^5^ MDA-MB-231 cells were seeded in 96-well plates for 12 h. Then MPCs were added and incubated for 6 h. For the PTT group, cells were treated with photothermal treatment for different action times (0, 4, 8, 12 and 16 min) and then placed back in the incubator for 6 h. For the MPST group, cells were treated with MMF for 2 h and then placed back in the incubator for 4 h. Then, the next steps for the MPST group were the same as those for the PTT group. Finally, the cells were then assayed for activity with the CCK-8 reagent. The synergistic effect index Q (Q ≥ 1.15: synergistic effect) was obtained by the following formula, where E_MMF_, E_PTT_ and E_MPST_ represent the respective treatment effects.

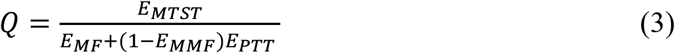

### 4.11. Cell Apoptosis Analysis

The Annexin V-FITC Apoptosis Detection Kit was used to assess the apoptosis of MDA-MB-231 cells. Cells (2 × 10^5^ pcs) were seeded in 6-well plates for 12 h to allow attachment according to the manufacturer’s instructions. Next, MPCs (300 μg/mL) were treated in different ways after coincubation with cells for 12 h. Afterwards, cells were collected and washed twice with PBS through centrifugation at 1000 g for 5 min, and then resuspended in binding buffer containing Annexin V-FITC and PI. After incubation in the dark for 15 min at room temperature, the cells were analyzed for apoptosis by flow cytometry (Fortessa, BD, America).

### 4.12. In Vivo Experiments of MDA-MB-231 Breast Cancer Model

Female BALB/c mice (6-8 weeks old) were purchased from Nanjing Cavins Biotechnology Co., Ltd. (Nanjing, China) and used according to the protocol approved by the Regional Ethics Committee for Animal Experimentation of Ningbo University, China (license number: SYXK (Zhe) 2019-0005). A subcutaneous mouse model carrying triple-negative breast cancer MDA-MB-231 was established by injecting 8 × 10^6^ MDA-MB-231 cells into the right abdomen of female nude mice. When the tumour volume reached 100 mm^3^, the mice were randomly divided into five groups (n ≥ 6) including PBS, MPCs, MMF, PTT and MPST. Then, the MPCs were injected into the nude mice through the tail vein on day 0, day 3 and day 6, respectively. Next, the corresponding experiments were performed on different groups after 8 h of ingestion, where the treatment parameters for MMF and PTT were 60 mT, 1.5 Hz, 2 h and 1.5 W/cm^2^, 15 min, respectively. The TUNEL staining was performed on the third day after treatment by randomly selecting one from each group. Additionally, body weight and tumour size were recorded every other day for 16 consecutive days. Then, mice were sacrificed, and organs and tumour tissues were collected and subjected to H&E staining and Prussian blue staining.

### 4.13. Statistical Analysis

All data were statistically analyzed using the software GraphPad Prism 7. Data for each group are reported as mean ± SD. For statistical significance, statistical analysis was performed using a t-test, \*\**p* < 0.01, \*\*\**p* < 0.001, \*\*\*\**p* < 0.0001.

## CRediT author statement

**Hui Du**: Conceptualization, Methodology, Investigation, Data curation, Writing – original draft, Visualization, Validation. **Fang Yang**: Conceptualization, Supervision, Writing – review & editing, Validation, Funding acquisition. **Chenyang Yao**: Methodology, Investigation. **Wenhao Lv**: Methodology, Visualization, Validation. **Hao Peng**: Methodology, Validation. **Stefan G. Stanciu**: Conceptualization, Validation. **Harald A. Stenmark**: Writing – review & editing, Validation. **Young Min Song**: Writing – review & editing. **Bo Jiang**: Validation. **Aiguo Wu**: Conceptualization, Project administration, Resources, Supervision, Funding acquisition. Hui Du and Yang Fang contributed equally to this work.

## Data availability

Data will be made available on request.

## Acknowledgements

This work is supported by the National Natural Science Foundation of China (32025021, 51803228, 31971292, 32111540257), the Youth Innovation Promotion Association, the Chinese Academy of Sciences (2022301), and the Ningbo 3315 Innovative Talent Project (2018-05-G). SGS and HAS acknowledge the support of the UEFISCDI grant RO-NO-2019-0601 MEDYCONAI. SGS, FY, and YMS acknowledge the support of the EsSENce CA19118 COST Action, which facilitated fruitful interactions.

## Appendix A. Supplementary data

Supplementary material related to this article can be found in the online version.

