## Supplementary for "“Double-Effect” strategy against Triple-Negative Breast Cancer *via* a synergistic therapy of Magneto-mechanical Force enhancing NIR-II Hypothermal Ablation"

### 1. Supplementary Methods

*1.1. Morphology and Structural Characterization of MPCs*

The morphology and size of MPCs were observed by TEM (HT7800, Hitachi) and HR-TEM (Talos F200X G2, Thermo Fisher). Inductively coupled plasma photoemission spectrometry (ICP-OES) was carried out using an Optima 2100 (PerkinElmer) system. X-ray diffraction (XRD, D8 Advance Davinci, Bruker) was used to determine the physical phase structure of MPCs. Dynamic Light Scattering (DLS, Zeta SIZER NANO ZS, Malvern Ltd.) was applied to measure hydrodynamic particle size and zeta potential. The Fourier Transform Infrared Spectrometer (FITR, Nicolet 6700, Thermo-fisher) was used to identify the functional groups attached to the surface of MPCs. In addition, UV-Vis-NIR spectrophotometers (Lambda 950, PerkinElmer) were utilized to assess the degree of light absorption by MPCs in the NIR region.

*1.2. Photothermal Effect*

The photothermal performance of the MPCs was evaluated through a series of experiments including the observation of the change in temperature rise for different concentrations of MPCs (0, 100, 200, 300, 400 μg/mL) irradiated with a 1064 nm laser (1.5 W/cm$^2$) for 10 min and a certain concentration of MPCs (300 μg/mL) irradiated with different laser power densities (0, 1.0, 1.5, 2.0 W/cm$^2$). In addition, the photothermal stability of MPCs (300 μg/mL) was tested for five on/off cycles under laser (1.5 W/cm$^2$) irradiation.

*1.3. Cell Culture*

The human breast cancer cell line (MDA-MB-231) was cultured in a DMEM medium containing 10% fetal bovine serum (FBS) in a humidified environment at 37 °C and 5% $CO_2$. When the cells grew to 80-90% confluence, these cells were digested and collected for different experimental evaluations. All cells were obtained from the Chinese Academy of Sciences Cell Bank (Shanghai, China).

*1.4. Cytotoxicity Measurement*

The biocompatibility of MPCs in the MDA-MB-231 cell line was assessed by using the CCK-8 assay kit. First, 100 μL of $10^5$/mL cells were seeded into 96-well plates and cultured for 12 h to adhere to the well, followed by co-incubation with different concentrations of MPCs (0, 50, 100, 200, 400, 600, 800 μg/mL) at 37 °C for 24 h. The cell viability was measured by using the CCK-8 assay kit under an enzyme calibrator (SpectraMax 190, MD) with an absorbance of 450 nm.

*1.5. Cellular Uptake and Localization*

(1) 300 μg/mL MPCs modified with MB dye were first co-incubated with cells ($1 \times 10^5$/mL) which had grown on confocal dishes for 12 h. After washing off the excess material with phosphate buffer solution (PBS) and fixing the cells with 1 mL of paraformaldehyde (PFA, 4%) for 30 min, the nuclei were stained with Hoechst. Finally, cells were observed by Confocal Laser Scanning Microscopy (CLSM, TCS SP8, Leica) with excitation of Hoechst and MB at 405 nm and 633 nm with emission ranges at 415-480 nm and 605-680 nm, respectively. (2) After co-incubation of 300 μg/mL of MPCs and cells for 12 h, the cells were collected by centrifugation (1000 rpm, 5 min) and fixed in 2.5% glutaraldehyde for 12 h at 4 °C. Afterwards, cells were dehydrated by acetone and embedded in Epon Araldite resin. Finally, the samples were sliced, stained, and observed under Bio-TEM at 80 kV.

*1.6. PTT Assessment In Vitro*

The photothermal effect of MPCs *in vitro* was assessed using the CCK-8 assay. Briefly, $1.2 \times 10^5$ cells were co-cultured with MPCs (300 μg/mL) in 96-well plates for 6 h and then washed three times with PBS and irradiated with a 1064 nm laser for different times (0, 4, 8, 12 and 16 min), accompanied by a thermal imager (JM500XC, Optris PI, Germany) with a PI Connect system to monitor the temperature rise. After that, the cells were placed back into the incubator and continued to incubate for 6 h. Finally, cell viability was assessed with CCK-8.

*1.7. Evaluation of ROS*

The DCFH-DA probe was used to detect the level of reactive oxygen species (ROS) inside the cancer cells after different modalities of treatment with MPCs. Briefly, cells ($1.2 \times 10^5$ pcs) were seeded in confocal discs for 12 h to adhere to the well, and then MPCs (300 μg/mL) were added for 12 h to coculture. After washing the cells three times with PBS and applying different treatments containing MF (60 mT, 15 Hz, 2 h), PTT (1.5 W/cm$^2$, 8 min) and MPST groups, they were stained with DCFH-DA for 20 min. In addition, cell nuclei were stained with Hoechst (1 μg/mL) for 10 min. Thereafter, the cell nuclei and ROS were imaged using a confocal laser scanning microscope (CLSM, TCS SP8, Leica) at excitation wavelengths of 488 nm and 515 nm, respectively.

#### *1.8. Calcein AM/Propidium Iodide Co-staining*

Oxidative damage to cancer cells by MPCs was investigated by Calcein AM/Propidium iodide (PI) co-staining. After co-culture with MPCs (300 μg/mL) for 6 h, cells were washed three times with PBS and treated in different ways, followed by staining with Calcein AM and PI for 30 min. Finally, cells were observed with an inverted fluorescence microscope (TS2R-FL, Nikon, Japan).

#### *1.9. Animal Safety Experiments*

To verify the biosafety of MPCs in animals, 100 μL of different concentrations of MPCs (0, 7.5, 15, 22.5 and 30 mg/Kg) were injected intravenously into ICR mice (n=7). After 14 days, blood was collected from each group of mice for blood routine and blood biochemical assays.

### 2. Supplementary Figures

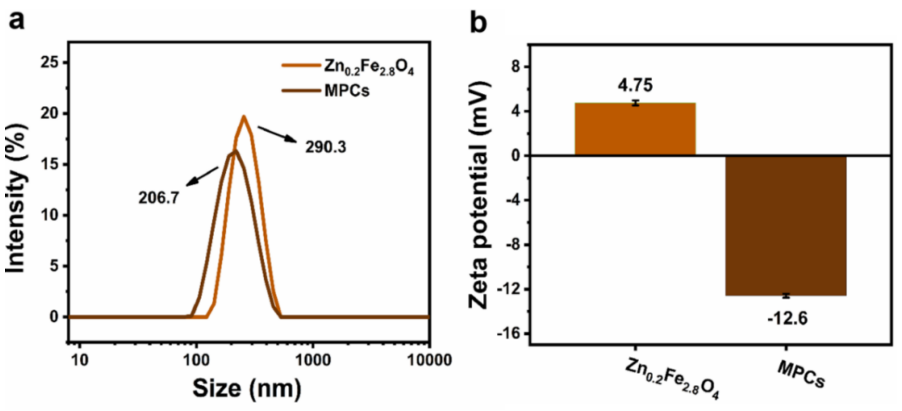

**Figure S1**. (a) The particle size distribution of $Zn_{0.2}Fe_{2.8}O_4$ NPs and MPCs was measured by dynamic light scattering, where the average diameter of MPCs was 206.7 nm with a PDI of 0.139, while that of $Zn_{0.2}Fe_{2.8}O_4$ was 290.3 and the PDI was 0.269. (b) Zeta potential of $Zn_{0.2}Fe_{2.8}O_4$ NPs and MPCs.

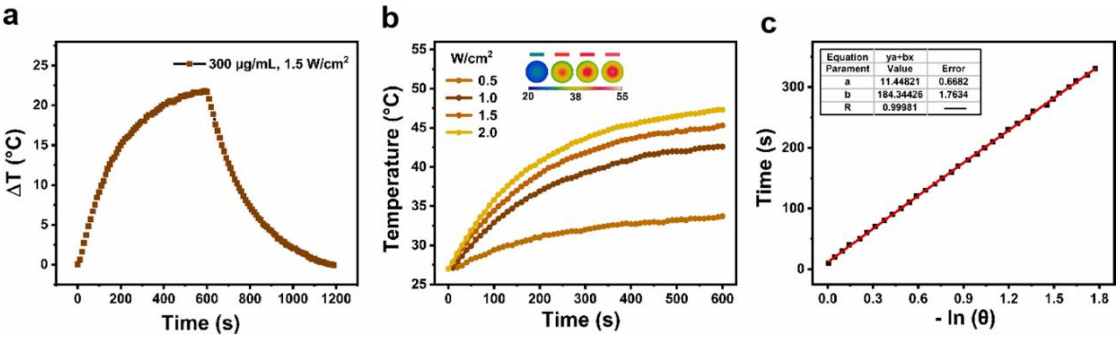

**Figure S2**. (a) Temperature rises of MPCs (300 μg/mL) under 1064 nm laser irradiation at 1.5 W/cm$^2$. (b) Photothermal conversion of MPCs (300 μg/mL) at different irradiation intensities of the 1064 nm laser. (c) Linear correlation between cooling time and the negative natural logarithm of the driving force temperature.

The photothermal conversion efficiency PCE (η) of MPCs was determined by irradiating 1064 (1.5 W/cm$^2$) for 8 min. Afterwards, the laser was turned off and the temperature was recorded. The value of PCE (η) was calculated according to the following equation reported in the literature [1].

$$\eta = [ hS (T_{max} - T_{surr}) - Q_{dis}] / I (1\text{-}10^{-A_{1064}}) \quad (1)$$

Where h is the heat transfer coefficient, S is the surface area, $T_{max}$ is the maximum temperature, and $T_{surr}$ refers to the surrounding temperature. The heat absorbed ($Q_{dis}$) can be neglected. I is the laser power intensity (1.5 W/cm$^2$) and the absorbance of MPCs is 0.86. The value of hS was calculated using the following equation:

$$hS = mD \times cD / \zeta s \quad (2)$$

$\zeta s$ is the heat dissipation time constant, which was identified by plotting linear data for the cooling period versus the negative natural logarithm, as shown below.

$$t = - \zeta s \ln(\theta) \quad (3)$$

$$\theta = [ T - T_{surr}] / [ T_{max} - T_{surr} ] \quad (4)$$

Thus， $hS= 0.3 \times 4.2 / 184.344$ J/sec °C = 6.84 mW/°C

$$\eta_{1064} = 6.84 \times 22 / 1500 (1 - 10^{-0.86}) = 11.65\%$$

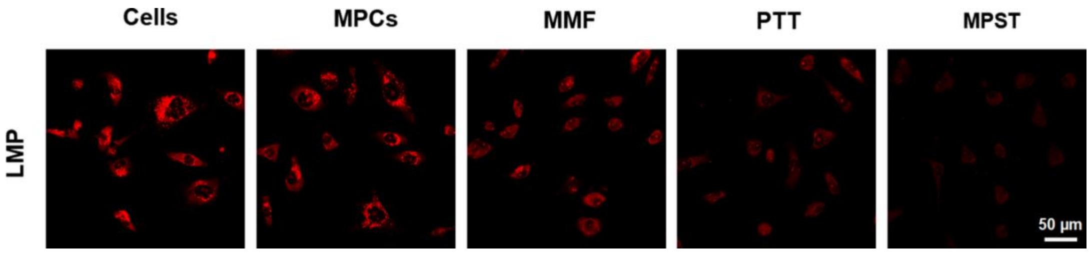

**Figure S3**. Lysosomal membrane permeability was characterized with the Acridine Orange Fluorescence Assay Kit. Representative images show that the decrease in the number of red fluorescent dots indicates an increase in LMP. Scale bar: 50 μm.

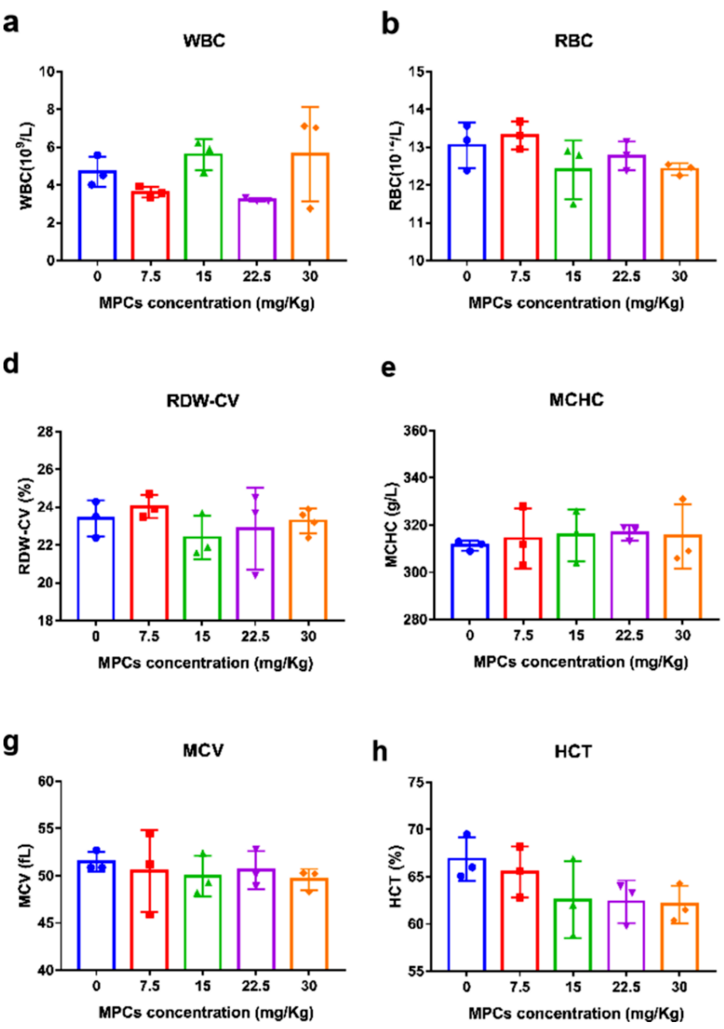

**Figure S4**. Hematological analysis of mice after uptake of different concentrations of MPCs.

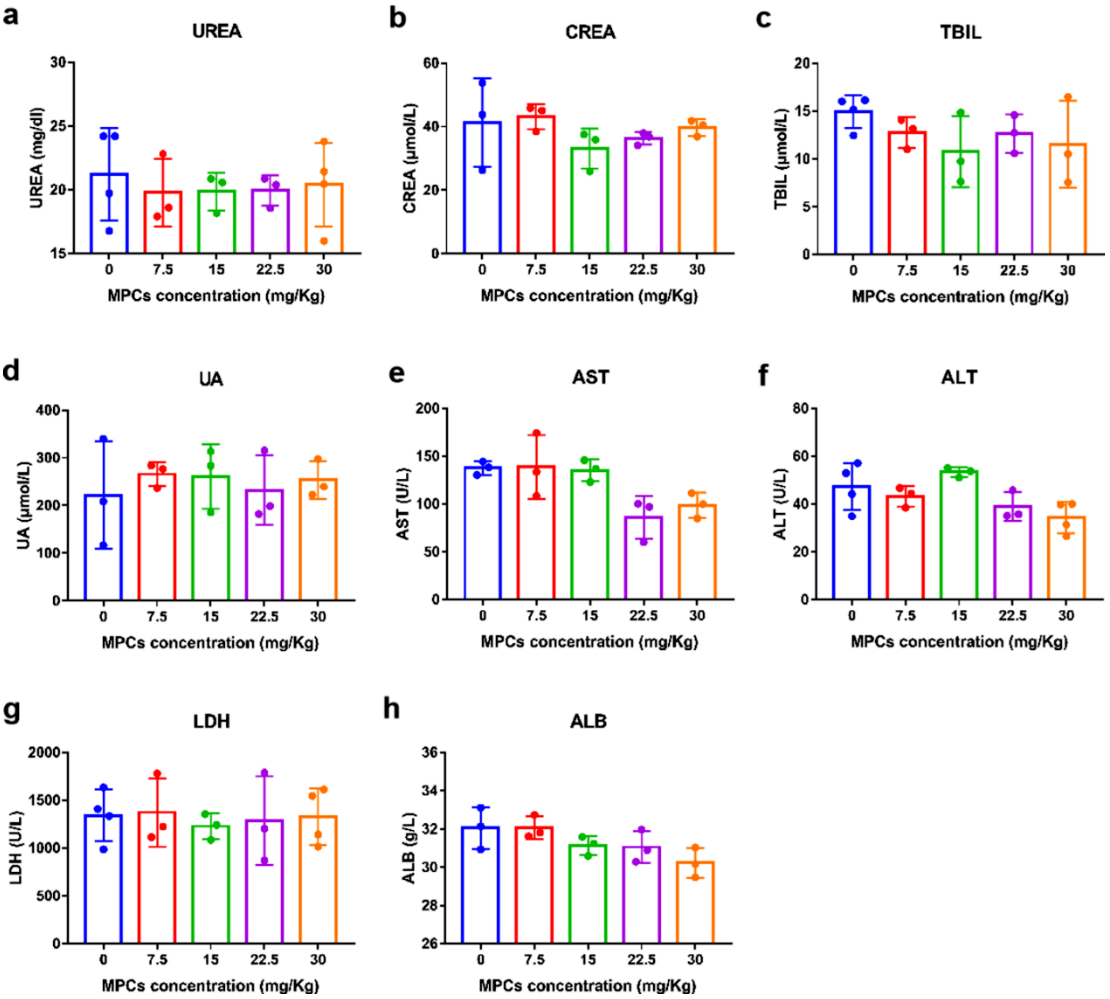

**Figure S5**. Biochemical analysis of mice blood after ingestion of various concentrations of MPCs.

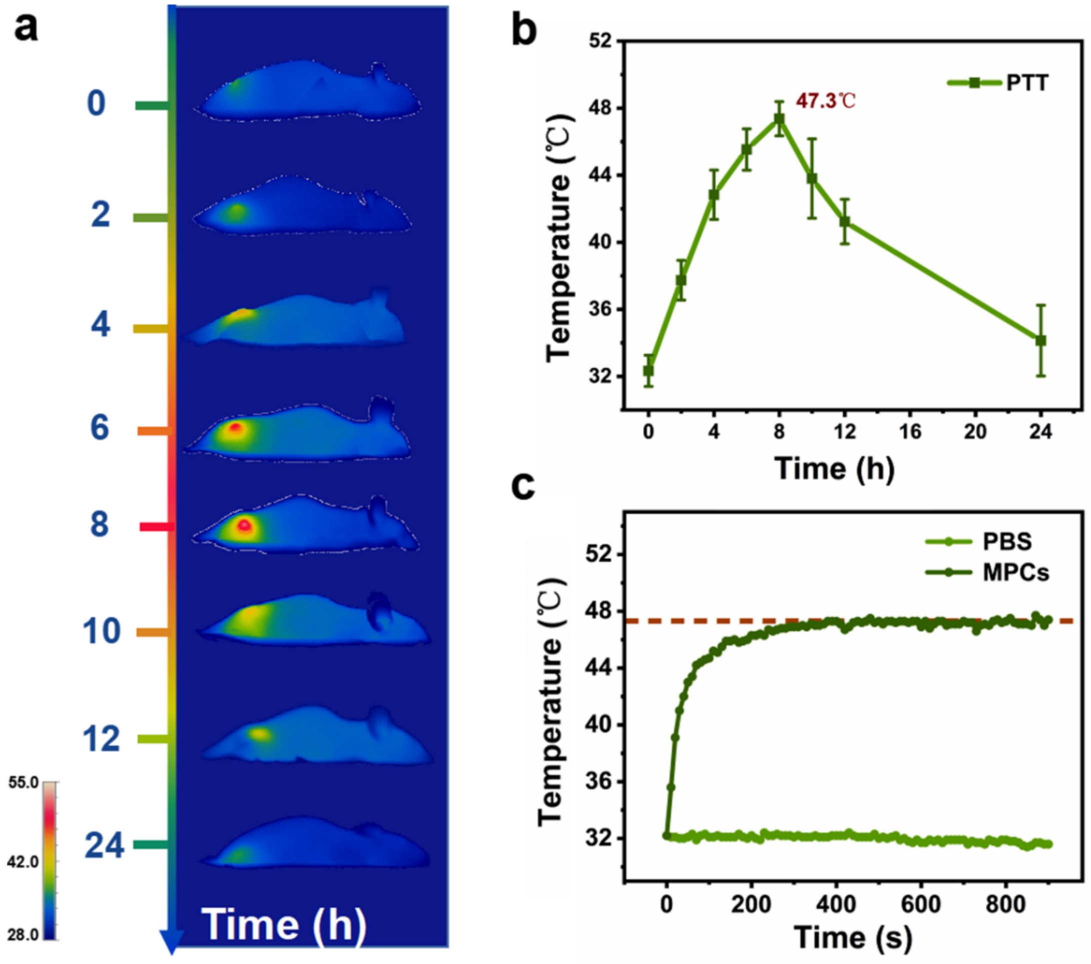

**Figure S6**. (a) Temperature rise in mice at different time points under 1064 nm laser irradiation after material uptake. (b) Quantification of the temperature rise of tumour sites (n=3) with localized MPCs at different periods under 1064 nm irradiation. (c) Temperature rise curve of MPCs in mice after 8 h intake at the tumour site by NIR light irradiation for 10 min.

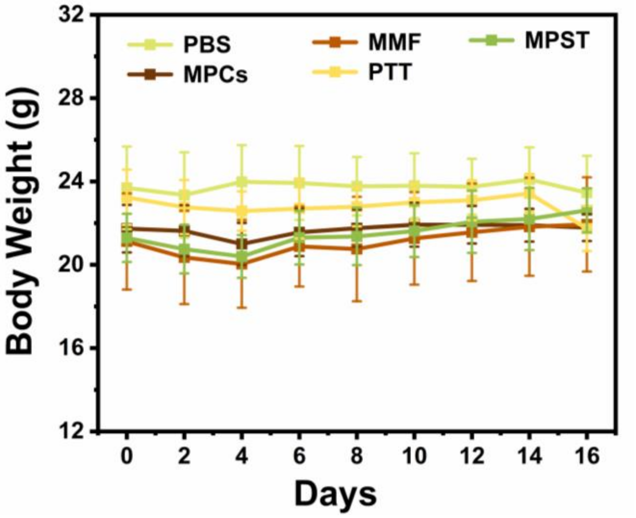

**Figure S7**. The changes in body weight of nude mice in 16 days under different treatments.

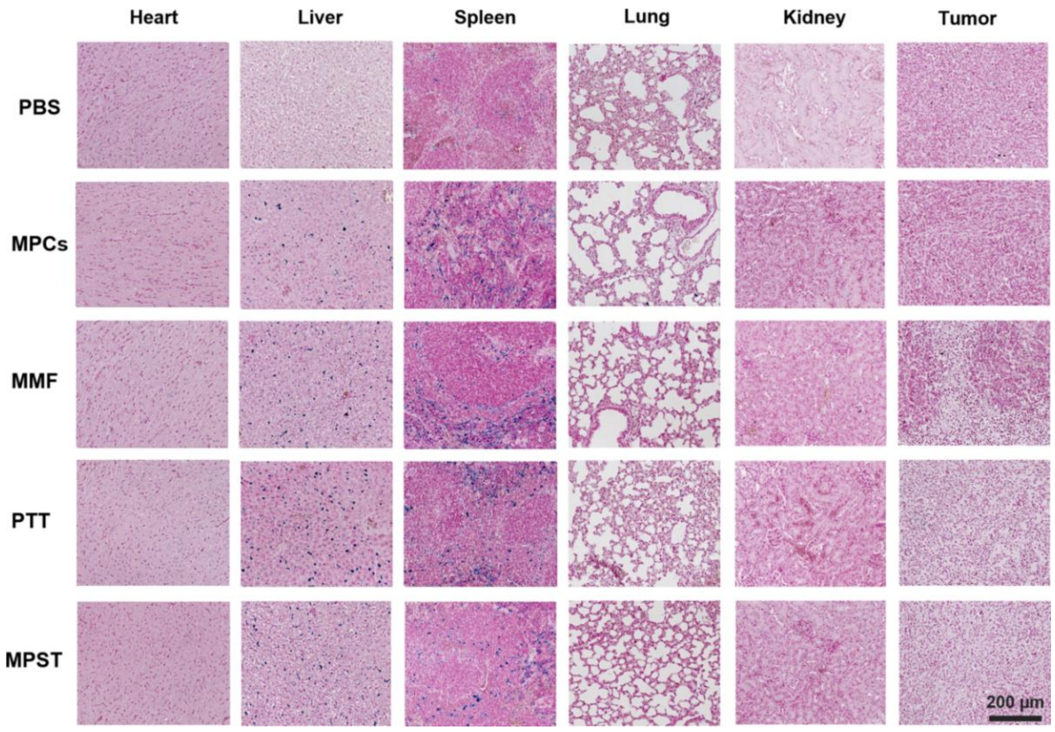

**Figure S8**. Prussian blue staining of tumours and organs was extracted from nude mice with MDA-MB-231 cells following 16 days of treatment (the scale bar: 200 μm).

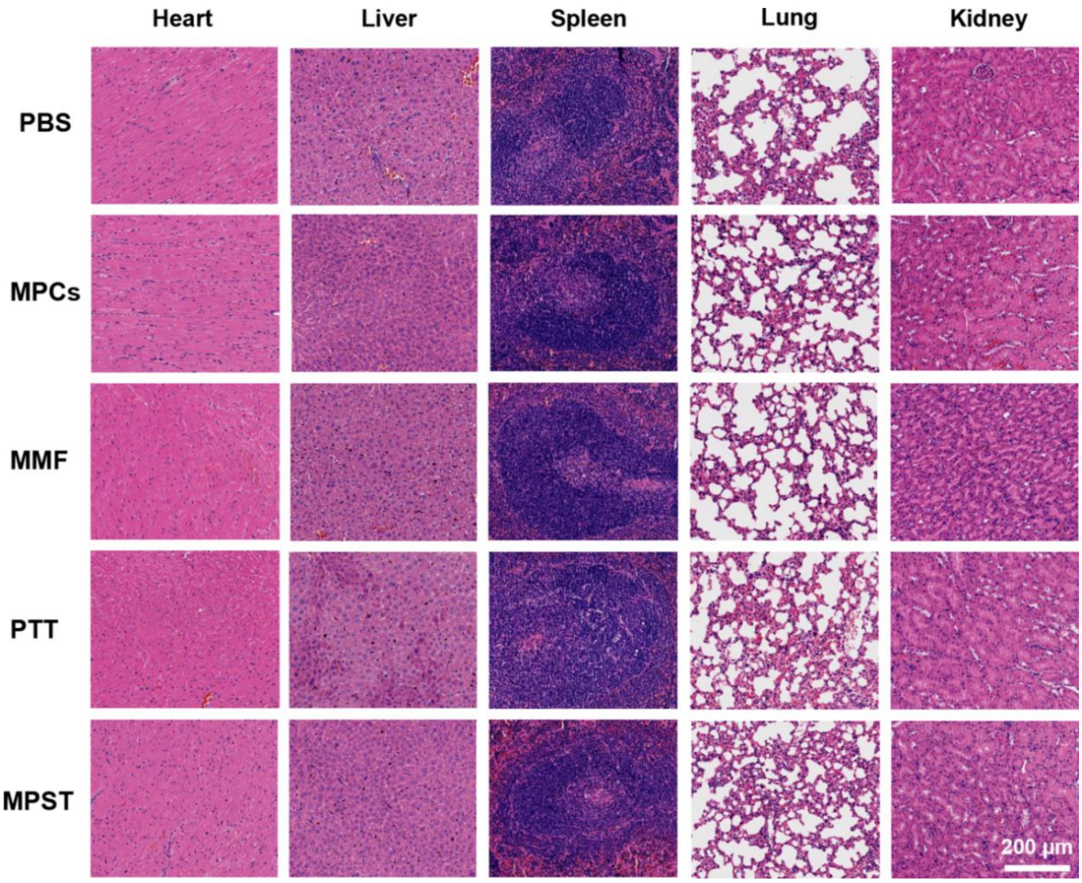

**Figure S9**. H&E staining of organs extracted out from nude mice bearing MDA-MB-231 cells following 16 days of treatment. No obvious lesion was found in normal tissue (scale bar: 200 μm).

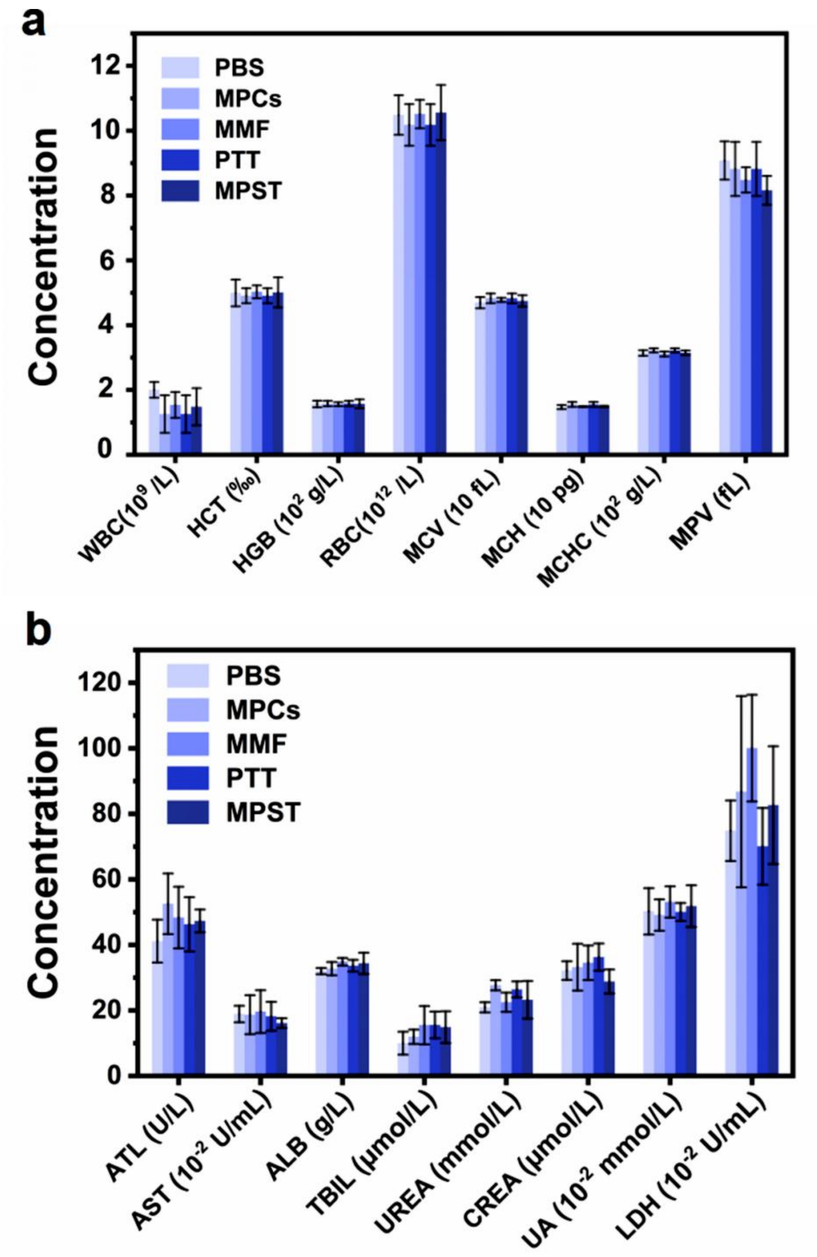

**Figure S10**. Hematological analysis (a) and blood biochemical analysis (b) of nude mice after different treatments (n=5).

#### 3. Supplementary Tables

**Table S1**. The real content of Zn doping in the $Zn_{0.2}Fe_{2.8}O_4$ MNPs by ICP-OES (Rr: real ratio of Fe and Zn content, Tr: theoretical ratio of Fe and Zn content)

| Sample | Fe (g/L) | Zn (g/L) | $C_{Fe}$ (M) | $C_{Zn}$ (M) | Rr | Tr | Sample |
|---|---|---|---|---|---|---|---|
| $Zn_{0.2}$ | 1.60 ±0.50 | 1.34 ±0.12 | 0.29 ±0.01 | 0.02 ±0.002 | 13.90 ±1.12 | 14 | $Zn_{0.2}Fe_{2.8}O_4$ |

**Table S2**. The ability of MMF to inhibit tumour cell migration

| Migration time (h) | Cell migration rate (%) | MMF migration rate (%) | MMF migration inhibition rate(%) |
|---|---|---|---|
| 12 | 16.88±0.35 | 8.37±1.43 | 50.42 |
| 24 | 31.41±2.51 | 12.22±1.50 | 61.10 |
| 36 | 54.87±1.3 | 17.76±2.67 | 67.63 |

**Table S3**. The ability of MMF to inhibit tumour cell invasion

| Invasion time (h) | Cell invasion number (pcs) | MMF invasion number (pcs) | MMFinvasion inhibition rate (%) |
|---|---|---|---|
| 36 | 896.5±109.02 | 252.67±70.59 | 71. 82 |

**Table S4**. MPST effects under different PTT durations

| PTT time (min) | PTT cell mortality (%) | RMFcell mortality (%) | MPST cell mortality (%) | Q value |
|---|---|---|---|---|
| 0 | 5.80±2.73 | 60 mT<br>15 Hz<br>2 h<br>25.19±1.56 | 30.02±1.93 | 1.02 |
| 4 | 8.61±2.97 | | 41.37±5.35 | 1.31 |
| 8 | 39.77±2.35 | | 76.59±0.92 | 1.39 |
| 12 | 44.37±1.73 | | 80.66±3.31 | 1.38 |
| 16 | 60.96±1.65 | | 91.48±2.28 | 1.29 |
